# Sex Chromosome Dosage Effects On Gene Expression In Humans

**DOI:** 10.1101/137752

**Authors:** Armin Raznahan, Neelroop Parikshak, Vijayendran Chandran, Jonathan Blumenthal, Liv Clasen, Aaron Alexander-Bloch, Andrew Zinn, Danny Wangsa, Jasen Wise, Declan Murphy, Patrick Bolton, Thomas Ried, Judith Ross, Jay Giedd, Daniel Geschwind

**Author notes:** **CORRESPONDING AUTHOR:** Armin Raznahan MD PhD, Developmental Genomics Unit, Rm 4D18, Building 10, 10 Center Drive, National Institute of Mental Health, NIH, Bethesda, MD, USA, E.

## Abstract

A fundamental question in the biology of sex-differences has eluded direct study in humans: how does sex chromosome dosage (SCD) shape genome function? To address this, we developed a systematic map of SCD effects on gene function by analyzing genome-wide expression data in humans with diverse sex chromosome aneuploidies (XO, XXX, XXY, XYY, XXYY). For sex chromosomes, we demonstrate a pattern of obligate dosage sensitivity amongst evolutionarily preserved X-Y homologs, and update prevailing theoretical models for SCD compensation by detecting X-linked genes whose expression increases with decreasing X- and/or Y-chromosome dosage. We further show that SCD-sensitive sex chromosome genes regulate specific co-expression networks of SCD-sensitive autosomal genes with critical cellular functions and a demonstrable potential to mediate previously documented SCD effects on disease. Our findings detail wide-ranging effects of SCD on genome function with implications for human phenotypic variation.

**SIGNIFICANCE STATEMENT:** Sex chromosome dosage (SCD) effects on human gene expression are central to the biology of sex differences and sex chromosome aneuploidy syndromes, but challenging to study given the co-segregation of SCD and gonadal status. We address this obstacle by systematically modelling SCD effects on genome wide expression data from a large and rare cohort of individuals with diverse SCDs (XO, XX, XXX, XXXX, XY, XXY, XYY, XXYY, XXXXY). Our findings update current models of sex chromosome biology by (i) pinpointing a core set of X- and Y-linked genes with “obligate” SCD sensitivity, (ii) discovering several non-canonical modes of X-chromosome dosage compensation, and (iii) dissecting complex regulatory effects of X-chromosome dosage on large autosomal gene networks with key roles in cellular functioning.

## INTRODUCTION

Disparity in SCD is fundamental to the biological definition of sex throughout much of the animal kingdom. In almost all eutherian mammals, females carry two X-chromosomes, while males carry an X- and a Y-chromosome: presence of the Y-linked SRY gene determines a testicular gonadal phenotype, while its absence allows development of ovaries (1). Sexual differentiation of the gonads leads to hormonal sex-differences that have traditionally been considered the major proximal cause for extra-gonadal phenotypic sex-differences. However, diverse studies, including recent work in transgenic mice that uncouple Y-chromosome and gonadal status, have revealed direct SCD effects on several sex-biased metabolic, immune and neurological phenotypes (2).

These findings - together with reports of widespread transcriptomic differences between pre-implantation XY and XX embryos (3, 4) - suggest that SCD has gene regulatory effects independently of gonadal status. However, genome-wide consequences of SCD remain poorly understood, especially in humans, where experimental dissociation of SCD and gonadal status is not possible. Understanding these regulatory effects is critical for clarifying the biological underpinnings of phenotypic sex-differences, and the clinical features of sex chromosome aneuploidy (5) [SCA, e.g. Turner (XO) and Klinefelter (XXY) syndrome], which can both manifest as altered risk for several common autoimmune and neurodevelopmental disorders (e.g. systemic lupus erythematosus and autism spectrum disorders) (6, 7). Here, we explore the genome wide consequences of SCD through comparative transcriptomic analyses amongst humans across a range of dosages including typical XX and XY karyotypes, as well as several rare SCA syndromes associated with 1, 3, 4 or 5 copies of the sex chromosomes. We harness these diverse karyotypes to dissect the architecture of dosage compensation amongst sex chromosome genes, and to systematically map the regulatory effects of SCD on autosomal gene expression in humans.

We examined gene expression profiles in a total of 470 lymphoblastoid cell lines (LCLs), from (i) a core sample of 68 participants (12 XO, 10 XX, 9 XXX, 10 XY, 8 XXY, 10 XYY, 9 XXYY) yielding for each sample genome-wide expression data for 19,984 autosomal and 894 sex-chromosome genes using the Illumina oligonucleotide Beadarray platform (**Methods**), and (ii) an independent set of validation/replication samples from 402 participants (4 XO, 146 XX, 22 XXX, 145 XY, 33 XXY, 16 XYY, 17 XXYY, 8 XXXY, 10 XXXXY) with quantitative reverse transcription polymerase chain reaction (qPCR) measures of expression for genes of interest identified in our core sample (**Table S1, Methods**).

## RESULTS

### Extreme Dosage Sensitivity of Evolutionarily Preserved X-Y Gametologs

To first verify our study design as a tool for probing SCD effects on gene expression, and to identify core SCD-sensitive genes, we screened all 20,878 genes in our microarray dataset to define which, if any, genes showed a persistent pattern of significant differential expression (DE) across all unique pairwise group contrasts involving a disparity in either X- or Y-chromosome dosage (n=15 and n=16 contrasts respectively, **Fig. 1a**). Disparities in X-chromosome dosage were always accompanied by statistically significant DE in 4 genes, which were all X-linked: *XIST* (the orchestrator of X-inactivation) and 3 other known genes known to escape X-chromosome inactivation (*PUDP*, *KDM6A*, *EIF1AX*) (8). Similarly, disparities in Y-chromosome dosage always led to statistically-significant DE in 6 genes, which were all Y-linked: *CYorf15B*, *DDX3Y*, *TMSB4Y*, *USP9Y*, *UTY*, and *ZFY*. Observed expression profiles for these 10 genes perfectly segregated all microarray samples by karyotype group (**Fig. 1b**), and could be robustly replicated and extended using available qPCR data for 5/5 of these genes in the independent sample of 402 LCLs from participants with varying SCD (**Fig. S1, Methods**).

**Figure 1.**
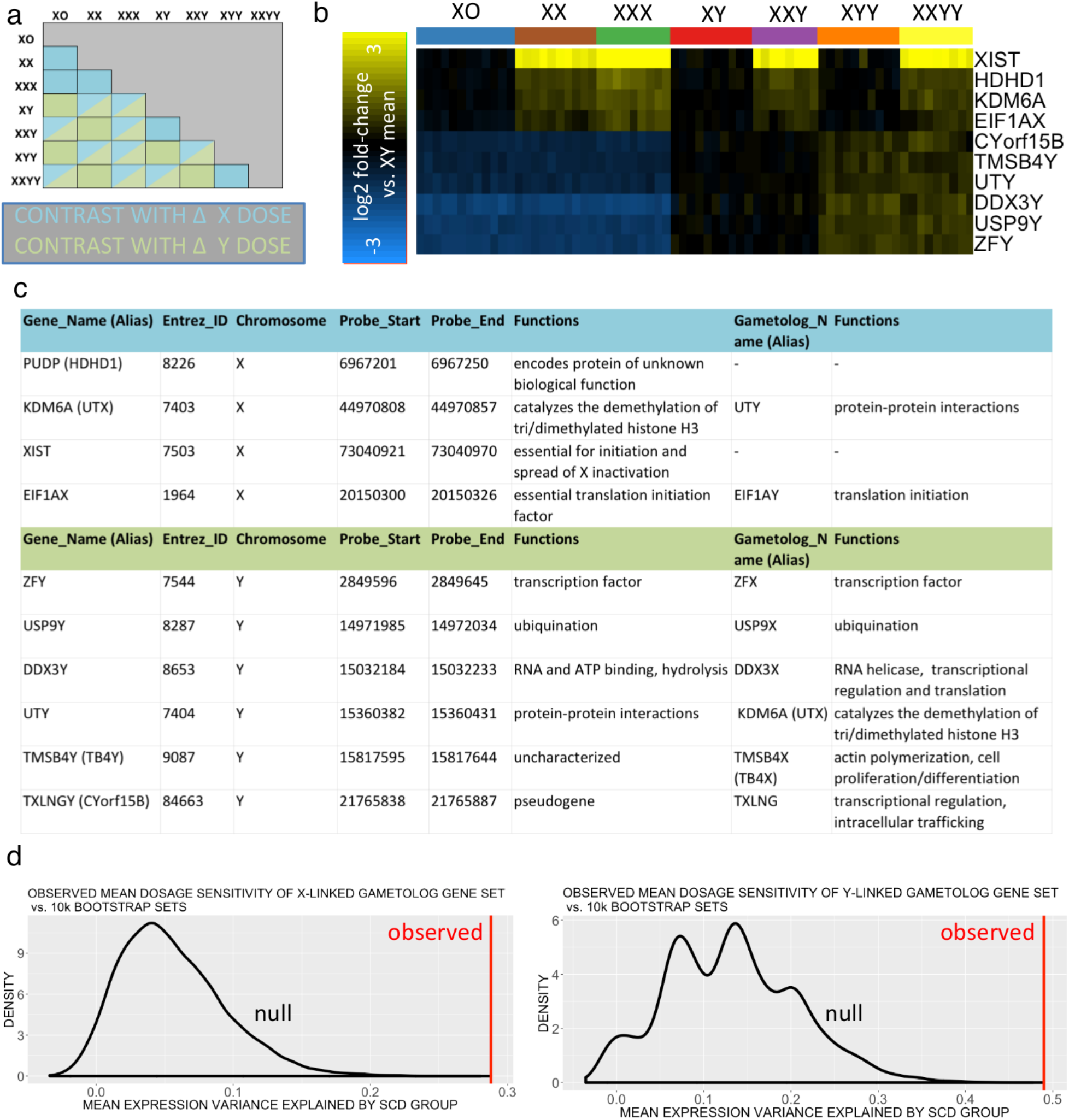
Consistent Gene Expression Changes with Altered Sex Chromosome Dosage. **a)** Cross table showing all unique pairwise SCD group contrasts within our microarray dataset, color coded for their involvement of changes in X- and Y-chromosome count. **b)** Two-dimensional expression heat-map for the 10 genes showing differential expression across all contrasts that involve disparity in X or Y-chromosome count. Column colors encode SCD group membership for each sample. Rows detail gene expression across all SCD samples as a log 2 fold change relative to the mean expression in XY males. **c)** Table providing Gene ID, location, function and homolog annotations for the 10 genes that showed obligate SCD sensitivity. Eight genes in this set are members of X-Y gametolog gene pairs. **d)** Density plots showing observed mean SCD sensitivity of the 14 gametolog genes in our study (red line), vs. null distribution (black line) of mean SCD sensitivity for 10,000 randomly sampled sets of non-gametolog sex-linked genes of equal size. Results are provided separately for X- and Y-chromosomes. For both chromosomes, the mean SCD sensitivity of the gametolog gene set is greater is than that of all 10k permuted gene sets.

Strikingly, 8 of the 10 genes showing obligatory SCD sensitivity (excepting XIST and PUPD) are members of a class of 16 sex-linked genes with homologs on both the X and Y chromosomes (i.e. 16 X-Y gene pairs, henceforth gametologs) (9) that are distinguished from other sex-linked genes by (i) their selective preservation in multiple species across ~300 million years of sex chromosome evolution to prevent male-female dosage disparity, (ii) the breadth of their tissue expression from both sex chromosomes; and (iii) their key regulatory roles in transcription and translation (9, 10) (**Fig. 1c**). Broadening our analysis to all 14 X-Y gametolog pairs present in our microarray data found that these genes as a group exhibit a heightened degree of SCD-sensitivity that distinguishes them from other sex-linked genes (**Fig. 1d, Methods**). These findings provide direct evidence that the evolutionary maintenance, broad tissue expressivity and enriched regulatory functions of X-Y gametologs (10) are indeed accompanied by a distinctive pattern of dosage sensitivity, which firmly establishes these genes as candidate regulators of SCD effects on wider genome function.

### Observed Sex Chromosome Dosage Effects on X- and Y-chromosome Genes Modify Current Models of Dosage Compensation

We next harnessed our study design to test the canonical (“Four Class”) model for SCD compensation, which defines four mutually-exclusive classes of sex chromosome genes that would be predicted to have differing responses to changing SCD(11): (i) pseudoautosomal region (PAR) genes, (ii) Y-linked genes, (iii) X-linked genes that undergo X-chromosome inactivation (XCI), and (iv) X-linked genes that “escape” XCI (XCIE). Under the Four Class Model, PAR genes would be predicted to increase their expression with increases in X- or Y-chromosome count, whereas expression of Y-linked genes would increase linearly with mounting Y-chromosome count. Due to the non-binary nature of gene silencing with XCI (12), theorized SCD effects on expression of XCI and XCIE genes represent extreme ends of an X-chromosome dosage sensitivity continuum: an X-linked genes that undergoes full silencing with XCI would show no expression change with changes in X-chromosome dosage, whereas an X-linked gene that undergoes complete escape from X-chromosome inactivation would show a linear increase in expression with increasing X-chromosome count.

To test this canonical “Four Class Model” we considered all sex chromosome genes and performed unsupervised *k*-means clustering of genes by their mean expression in each of the 7 karyotype groups represented in our microarray dataset, and compared this empirically-defined grouping with that given *a priori* by the Four Class Model (**Methods**). *k*-means clustering distinguished 5 reproducible (color-coded) clusters of SCD-sensitive sex-chromosome genes that overlapped strongly with gene groups predicted by the Four Class Model, from a large left-over cluster of 773 genes with low or undetectable expression levels in most samples (median detection rate of 4/68 samples), and no significant SCD sensitivity (**Table S2, Fig. 2a,b, Fig. S2a,b**).

**Figure 2.**
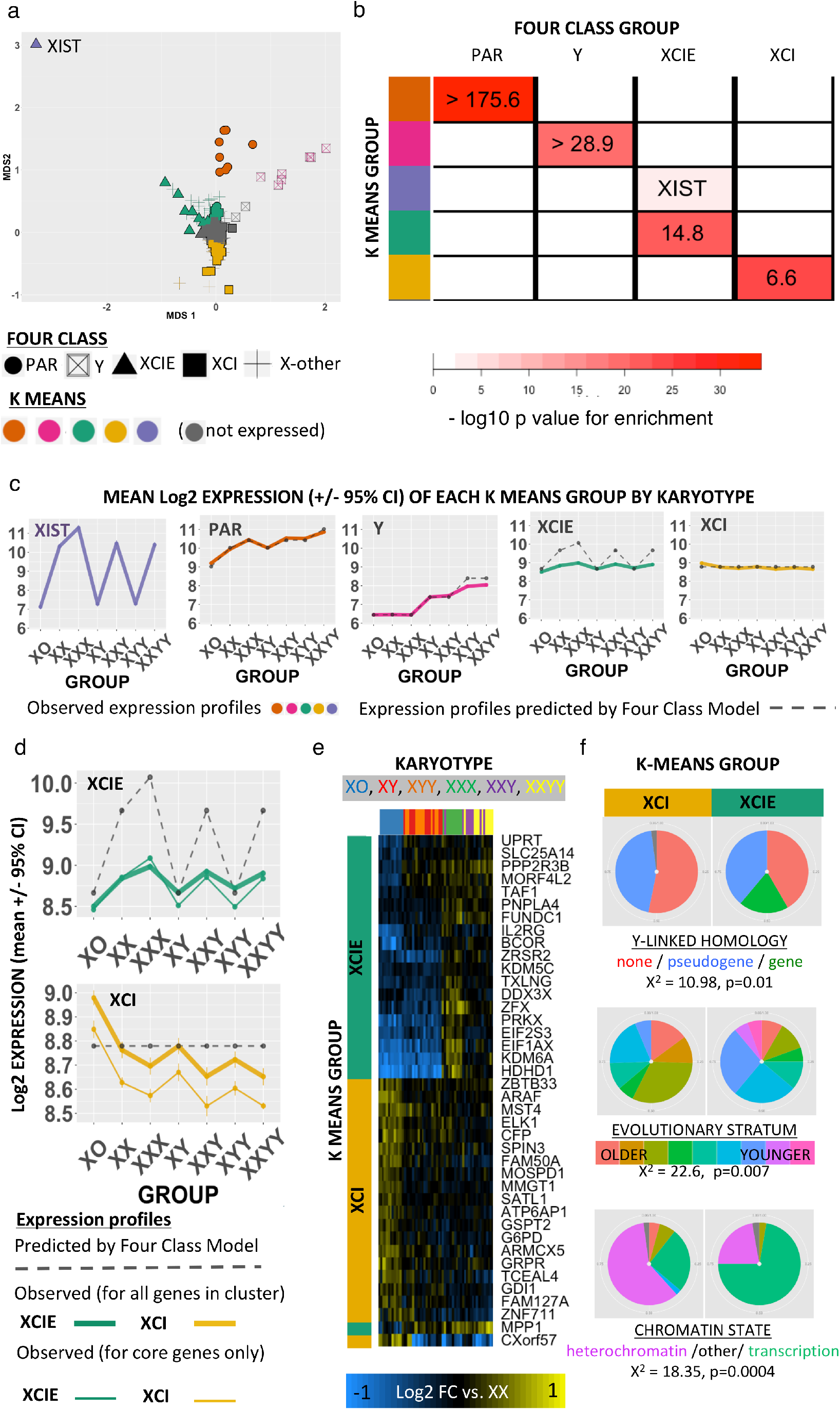
Data-Driven Partitioning of Sex Chromosome Genes by Dosage Sensitivity. **a)** 2D Multidimensional scaling (MDS) plot of sex chromosome genes by their mean expression profiles (+/- 95% confidence intervals) across all 7 SCD groups. Genes are coded by both Four Class Model and k-means cluster grouping. Note that MDS2 arranges X-linked genes along the established gradient of X-linked dosage sensitivity that ranges from extreme XCIE (XIST), to full XCI. **b)** Cross table showing enrichment of k-means clusters (rows) for Four Class Model gene groups. Lower-bounds for enrichment odds-ratios are given where mean enrichment = ∞. **c)** Dot and line plots showing observed and predicted mean expression values for each k-means gene cluster across karyotype groups. **d)** Close-up of observed (solid, color-coded) vs. predicted (dashed, gray) mean (+/- 95% confidence intervals) expression profiles of Green and Yellow gene clusters. Observed expression profiles still counter predictions when analysis is restricted to core genes in each cluster with XCIE/XCI status that has been confirmed across three independent reports (Balaton et al). **e)** Heatmap showing normalized (vs. XX mean) expression of dosage sensitive genes in the XCIE and XCI k-means groups (rows, color-coded green and yellow respectively), for each sample (columns, color coded by SCD group). **f)** Pie-charts showing how genes within XCIE and XCI-enriched k-means clusters (green and yellow columns respectively), display mirrored over/under-representation for three genomic features of X-linked gene that have been linked to XCEI in prior research (i) persistence of a surviving Y-linked homolog, (ii) location of the gene within “younger” evolutionary strata of the X-chromosome, and (iii) presence of euchromatic rather than heterochromatic epigenetic markers.

The 5 SCD-sensitive groups of sex chromosome genes detected by k-means were highly reproducible over k-means analyses across 1000 bootstrap draws from our sample pool (**Fig. S2b**), and consisted of: an Orange cluster of PAR genes, a Pink cluster of Y-linked genes (especially enriched for Y gametologs, odds ratio=5213, p=1.3*10^−15^), a Green cluster enriched for known XCIE genes (especially X gametologs, odds ratio=335, p=3.4*10^−11^), and a Yellow cluster enriched for known XCI genes. The X-linked gene responsible for initiating X-inactivation - XIST - fell into its own Purple “cluster” (**Fig. 2b**). For all but the Orange cluster of PAR genes, observed patterns of gene-cluster dosage sensitivity across karyotype groups deviated from those predicted by the Four Class Model (**Fig. 2c**).

Mean Expression for the Pink cluster of Y-linked genes increased in a stepwise fashion with Y-chromosome dosage, but deviated from the Four Class Model prediction by showing a sub-linear relationship with Y-chromosome count – indicting that these Y-linked genes may be subject to active dosage compensation. Fold-changes observed by microarray for 3/3 of these Y-linked genes were highly correlated across group contrasts with fold-changes observed between karyotype groups by qPCR in an independent sample of 402 participants with varying SCD (**Methods, Fig. S2d).**

Observed expression profiles the Yellow and Green clusters of X-linked genes also deviated from predictions of the Four Class Model predictions (**Table S2, Fig. 2c,d**). Linear models for X- and Y-chromosome dosage effects on expression (**Methods**) indicated that the XCI-enriched Yellow cluster was highly sensitive to SCD (F=47.7, p<2.2*10^−16^), and that the expression of this cluster was significantly *inversely* related to X-chromosome dosage at the level of both mean cluster expression (coefficient for linear effect of X-chromosome count on expression = -0.12, p=3.8*10^−15^) and the individual expression profile of 60/66 genes within the cluster (p<0.05 for negative linear effect of X count on expression). This observation suggests that increasing X copy number may not solely involve silencing of these genes from the additional inactive X-chromosome, but a further repression of their expression from the single active X-chromosome.

Remarkably, expression of the XCI gene cluster was also significantly decreased by presence of a Y-chromosome, at the level of both mean cluster expression (coefficient for linear effect of Y-chromosome count on expression = -0.09, p= 1.4*10^−14^) and expression profiles of 48/66 individual cluster genes (p<0.05 for negative linear effect of Y count on expression). The Green XCIE cluster manifested an inverted version of this effect whereby increases in Y chromosome dosage were associated with increased gene expression (p<6.2*10^−11^ for mean cluster expression and p<0.05 for 23/39 cluster genes) - providing the first evidence that Y-chromosome status can influence the expression level of X-linked genes independently of circulating gonadal factors. Mean expression of Green XCIE cluster genes also scaled sub-linearly with X-chromosome dosage. For both Green and Yellow clusters, we established that observed patterns of dosage sensitivity held when analysis was restricted to X-linked genes with only high confidence annotations for XCIE and XCI status (respectively) (**Fig 2d**) – suggesting that observed expression profiles was unlikely to be explained by misclassification of X-linked genes by XCI status. To further probe the sublinear relationship between XCIE Green cluster expression and X-chromosome dosage, we integrated our findings with those of a recently-published (13) survey of allelic expression imbalance analyses from female LCLs with skewed X-inactivation (**Text S1, Fig. S2**). This analysis revealed that the magnitude of observed sub-linear relationships between Green cluster gene expression values and X-chromosome dosage is consistent with independent measurements of incomplete of escape from XCI (12, 13).

To determine the reproducibility and convergent validity of the unexpected modes of dosage sensitivity observed for XCI (reduced expression with increasing X- and Y-chromosome dosage) and XCIE (increased expression with increasing Y-chromosome dosage) clusters, we first confirmed that the distinct expression profiles for these two clusters were reproducible at the level of individual genes and samples. Indeed, unsupervised clustering of microarray samples based on expression of XCI and XCIE cluster genes relative to XX controls distinguished three broad karyotype groups: females with one X-chromosome (XO), males with one X-chromosome (XY, XYY), and individuals with extra X-chromosome (XXX, XXY, XXYY) (**Fig. 2e**). We were also able to validate our data-driven discovery of XCI and XCIE gene clusters against independently generated X-chromosome annotations (**Fig. 2f**), which detail 3 distinct genomic predictors of inactivation status for X-linked genes. Specifically, XCI cluster genes were relatively enriched (and XCIE cluster genes relatively impoverished) for (i) having lost a Y-chromosome homolog during evolution (14) (X^2 =^ 10.9, p=0.01), (ii) being located in older evolutionary strata of the X-chromosome (15) (X^2 =^ 22.6, p=0.007), and (iii) bearing heterochromatic markers (16) (X^2 =^ 18.35, p=0.0004).

Finally, qPCR assays in LCLs from an independent sample of 402 participants with varying SCD validated the fold changes observed in microarray data for 5/6 of the most SCD-sensitive XCIE and XCI cluster genes (**Methods, Fig. S2e**). To independently extend these observations, we measured gene expression by qPCR in novel karyotype groups not represented in our microarray dataset (XXXY, XXXXY, **Methods**) and were able to confirm reduction in expression with greater X-chromosome dosage for 2 of 3 XCI cluster genes (**Fig S2f**, *NGFRAP1*, *CXorf57*), and Y-chromosome dosage effects upon expression for 5 of 6 X-linked genes from XCI and XCIE clusters (**Fig S2g**, downregulation: *NGFRAP1*, *CXorf57* | up-regulation: *PIM2*, *PRKX*). Taken together, these findings update the canonical Four Class Model of SCD compensation for specific Y-linked and X-linked genes, and expand the list of X-linked genes capable of mediating wider phenotypic consequences of SCD variation.

### Context-Specific Disruption of Autosomal Expression by Sex Chromosome Aneuploidy

We next leveraged the diverse SCAs represented in our study to assess how SCD variation shapes expression on a genome-wide scale. By counting the total number of differentially expressed genes (DEGs, **Methods**) in each SCA group relative to its respective euploidic control (i.e XO and XXX compared with XX; XXY, XYY, XXYY compared with XY), we detected order of magnitude differences in DEG count amongst SCAs across a range of log2 fold change (log2FC) cut-offs (**Fig. 3a,b**). We observed an order of magnitude increase in DEG count with X-chromosome supernumeracy in males vs. females, which although previously un-described, is congruent with the more severe phenotypic consequences of X-supernumeracy in males vs. females (17). Overall, increasing the dosage of the sex chromosome associated with the sex of an individual (i.e. X in females and Y in males) had a far smaller effect than other types of SCD changes. Moreover, the ~20 DEGs seen in XXX contrasted with >2000 DEGs in XO – revealing a profoundly asymmetric impact of X-chromosome loss vs. gain on the transcriptome of female LCLs, which echoes the asymmetric phenotypic severity of X-chromosome loss (Turner) vs. gain (XXX) syndromes in females (6).

**Figure 3.**
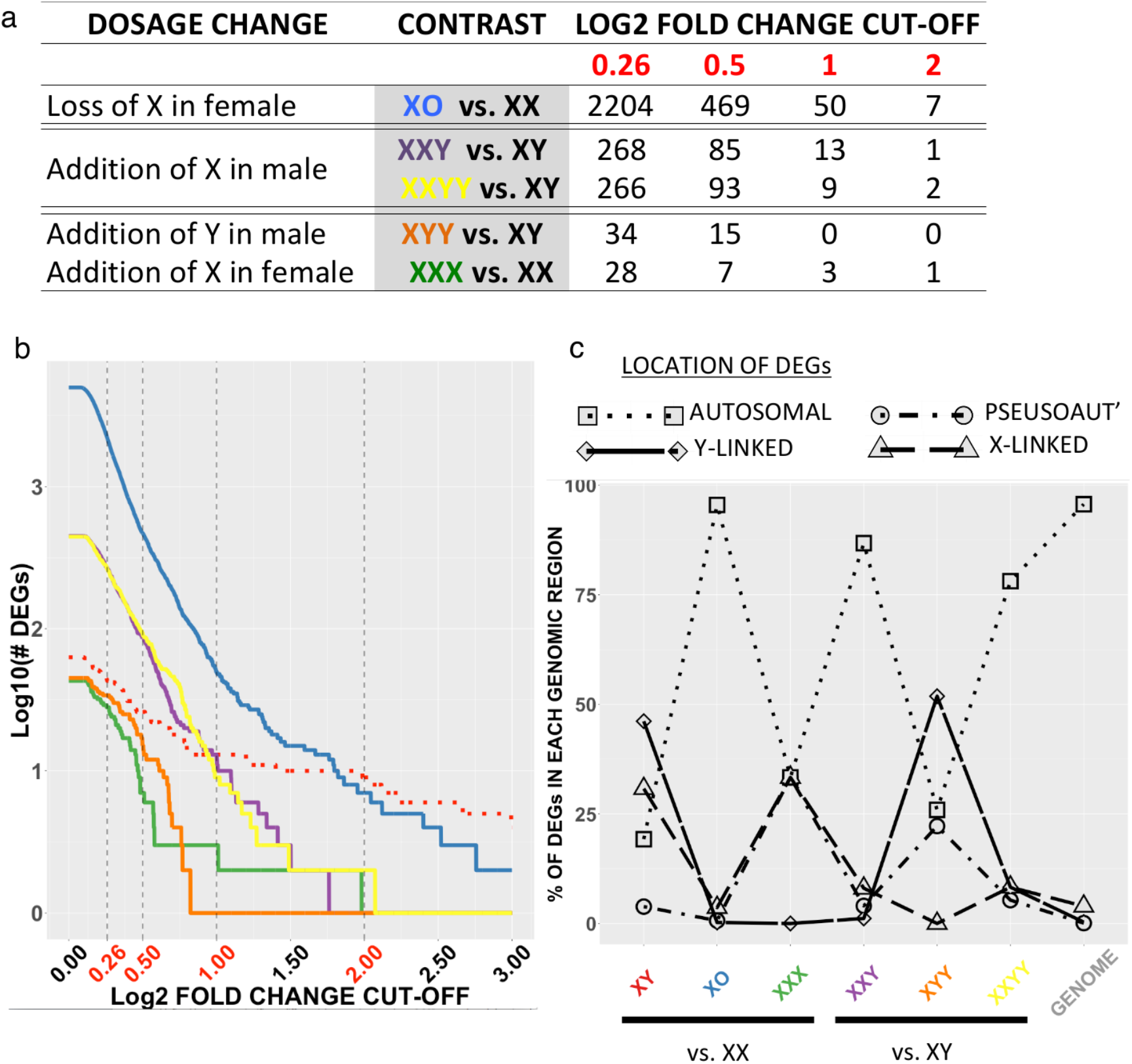
Genome-wide Effects of Sex Chromosome Dosage Variation. **a-b)** Table **a** and corresponding line-plot **b** showing number of genes with significant differential expression (after FDR correction with q<0.05) in different SCD contrasts at varying |log2 fold change| cut-offs. Note the order-of-magnitude differences between the number of Differentially Expressed Genes (DEGs) in XO (“removal of X from female”) vs. XXY and XXYY (“addition of X to male”) vs. XYY and XXX (“addition of Y and X to male and female, respectively”). A |log2 fold change| threshold of 0.26 (~20% change in expression) was applied to categorically define differential expression in other analyses, by identifying the log2 fold change threshold increase causing the greatest drop in DEG count for each karyotype group, and then averaging this value across karyotype groups. **c)** Dot-and-line plot showing the proportion of DEGs in each karyotype group that fell within different regions of the genome. The proportion of all genes in the genome within each genomic region is shown for comparison. All SCD groups showed non-random DEG distribution relative to the genome (p<2*10^-16), but DEG distributions differed significantly between SCD groups (p<2*10^-16). XO, XXX and XXYY are distinguished from all other SCDs examined by the large fraction of their overall DEG count that comes from autosomal genes.

To clarify the relative contribution of sex chromosome vs. autosomal genes to observed DEG counts with changes in SCD, we calculated the proportion of DEGs in every SCD group (comparing SCAs to their “gonadal controls”, and XY males to XX females) that fell within each of four distinct genomic regions: autosomal, PAR, Y-linked and X-linked (**Fig. 3c**). Autosomal genes accounted for >75% of all DEGs in females with X-monosomy (XO) and males with X-supernumeracy (XXY, XXYY), but <30% DEGs in all other SCD groups (**Methods**). These results reveal that SCD changes vary widely in their capacity to disrupt genome function, and demonstrate that differential involvement of autosomal genes is central to this variation. Moreover, associated SCA differences in overall DEG count broadly recapitulate SCA differences in phenotypic severity.

### Sex Chromosome Dosage Regulates Large-Scale Gene Co-expression Networks

To provide a more comprehensive systems-level perspective on the impact of SCD on genome-wide expression patterns, we leveraged Weighted Gene Co-expression Network Analysis (18) (WGCNA, **Methods**). This analytic approach uses the correlational architecture of gene expression across a set of samples to detect sets (modules) of co-expressed genes. Using WGCNA, we identified 18 independent gene co-expression modules in our dataset (**Table S3**). We established that these modules were not artifacts of co-differential expression of genes between groups by demonstrating their robustness to removal of all group effects on gene expression by regression (**Fig. S3a**), and after specific exclusion of XO samples (**Fig. S3b**) given the extreme pattern of DE in this karyotype (**Fig. 3b**). We focused further analysis on modules meeting 2 independent statistical criteria after correction for multiple comparisons: (i) significant omnibus effect of SCD group on expression, (ii) significant enrichment for one or more gene ontology (GO) process/function terms (**Methods, Table S3, Fig. 4a-b**). These steps defined 8 functionally coherent and SCD-sensitive modules (Blue, Brown, Green, Purple, Red, Salmon, Tan and Turquoise). Notably, the SCD effects we observed on genome wide expression patterns appeared to be specific to shifts in sex chromosome gene dosage, as application of our analytic workflow to publically available genome-wide Illumina beadarray expression data from LCLs in patients with trisomy 21 (Down syndrome) revealed a highly dissimilar profile of genome-wide expression change to that observed in sex chromosome trisomies (**Methods, Fig. 4c, Table S4**).

**Figure 4.**
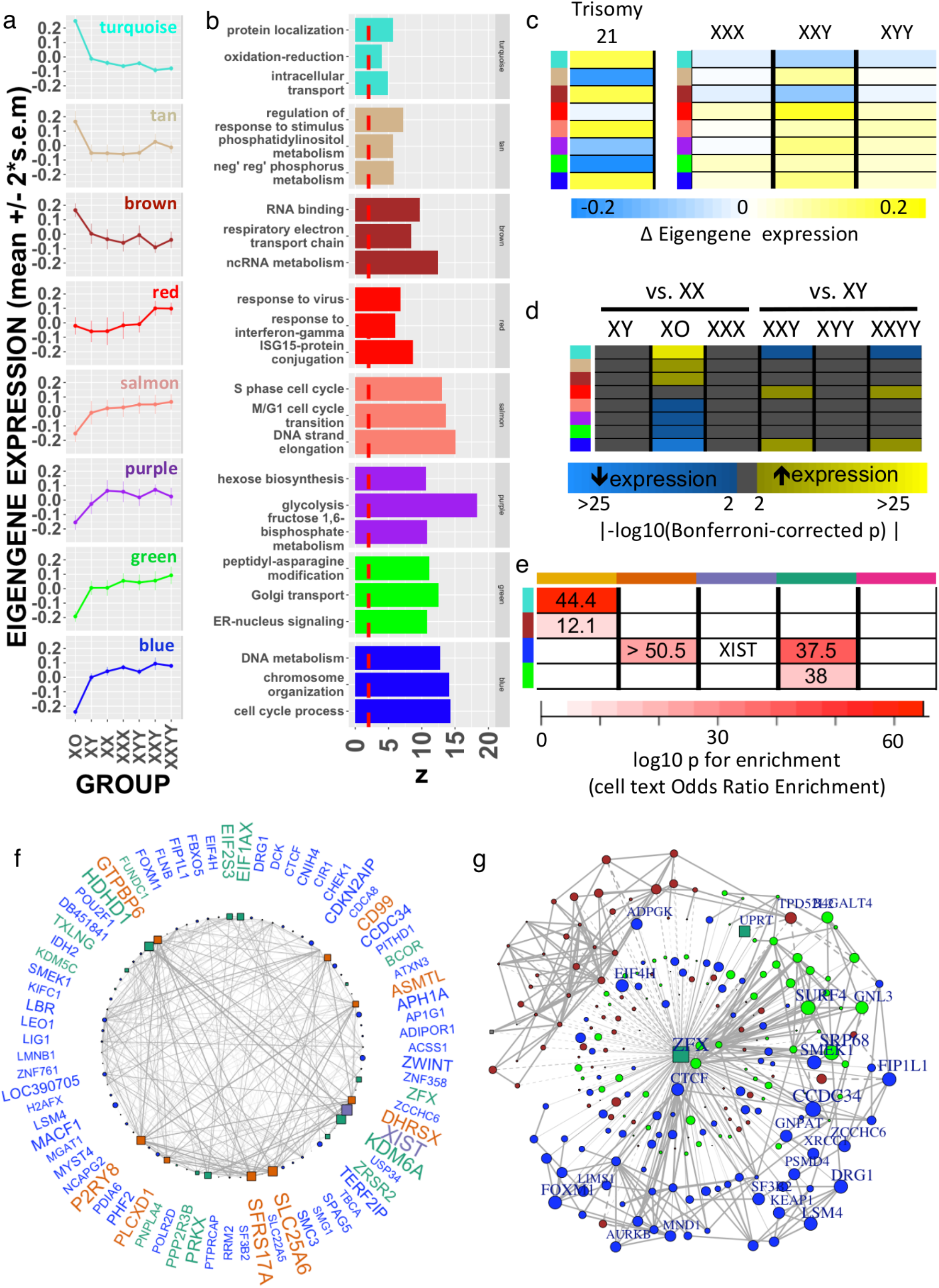
Weighted Gene Co-expression Network Analysis of Sex Chromosome Dosage Effects. **a)** Dot and line plots detailing mean expression (+/- 95% confidence intervals) by SCD group for 8 SCD-sensitive and functionally-coherent gene co-expression modules. **b)** Top 3 GO term enrichments for each module. **c)** Heatmap showing distinct profile of module DE with a supernumerary chromosome 21 vs. a supernumerary X-chromosome. **d)** Heatmap showing statistically significant differential expression of gene co-expression modules between karyotype groups. **e)** Cross tabulation showing enrichment of each module for the dosage-sensitive clusters of sex-chromosome genes detected by k-means. Lower-bounds for enrichment odds-ratios are given where mean enrichment = ∞. **f)** Gene co-expression network for the Blue module showing the top decile of co-expression relationships (edges) between the top decile of SCD-sensitive genes (nodes). Nodes are positioned in a circle for ease of visualization. Node shape distinguishes autosomal (circle) from sex chromosome (square) genes. Sex chromosome genes within the blue module are color coded by their SCD-sensitivity grouping as per **Figure 2a** (PAR-Orange, XCIE – dark green, XIST–purple). Larger node and gene name sizes reflect greater SCD sensitivity. Edge width indexes the strength of co-expression between gene pairs. **g).** ZFX and its target genes from Blue, Green and Brown modules with significant ZFX TFBS enrichment. Note that expression levels of ZFX (which increases in expression with mounting X-chromosome dosage) are positively correlated (solid edges) with SCD sensitive genes that are up-regulated by increasing X-chromosome dose (Blue and Green modules), but negatively correlated (dashed edges) with genes that are down-regulated by increasing X-chromosome dose (Brown module).

To specify SCA effects on module expression, we compared all aneuploidy groups to their respective “gonadal controls” (**Fig. 4d**). Statistically significant differences in modular eigengene expression were seen in XO, XXY and XXYY groups - consistent with these karyotypes causing larger total DE gene counts than other SCD variations (**Fig. 3a**). The largest shifts in module expression were seen in XO, and included robust up-regulation of protein trafficking (Turquoise), metabolism of non-coding RNA and mitochondrial ATP synthesis (Brown), and programmed cell death (Tan) modules, alongside down-regulation of cell cycle progression, DNA replication/chromatin organization (Blue, Salmon), glycolysis (Purple) and responses to endoplasmic reticular stress (Green) modules. Module DE in those with supernumerary X chromosomes on an XY background, XXY and XXYY, involved “mirroring” of some XO effects – i.e. *down-regulation* of protein trafficking (Turquoise) and *up-regulation* of cell-cycle progression (Blue) modules – plus a more karyotype-specific up-regulation of immune response pathways (Red).

The distinctive up-regulation of immune-system genes in samples of lymphoid tissue from males carrying a supernumerary X-chromosome carries potential clinical relevance for one of the best-established clinical phenotypes in XXY and XXYY syndromes: a strongly (up to 18-fold) elevated risk for autoimmune disorders (ADs) such as Systemic Lupus Erythmatosus, Sjogren Syndrome, and Diabetes Mellitus (7). In further support of this interpretation, we found the Red module to be significantly enriched (p=0.01 by Fisher’s Test, and p=0.01 by gene set permutation) for a set of known AD risks compiled from multiple large-scale Genome Wide Association Studies (GWAS, **Methods**). The two GWAS implicated AD risk genes showing strongest connectivity within the Red module and up-regulation in males bearing an extra X-chromosome were *CLECL1* and *ELF1* – indicating that these two genes should be prioritized for further study in mechanisms of risk for heightened autoimmunity in XXY and XXYY males. Collectively, these results represent the first systems-level characterization of SCD effects on genome function, and provide convergent evidence that increased risk for AD risk in XXY and XXYY syndrome may be arise due to an up-regulation of immune pathways by supernumerary X-chromosomes in male lymphoid cells.

To test for evidence of coordination between the changes in sex-chromosome genes imparted by SCD (**Fig. 2**), and the genome-wide transcriptomic variations detected through WGCNA (**Fig. 4a**), we asked if any SCD-sensitive gene co-expression modules were enriched for one or more of the 5 SCD-sensitive clusters of sex chromosome genes (i.e. “PAR”, “Y-linked”, “XCIE”, “XCI” and the gene XIST). Four WGCNA modules - all composed of >95% autosomal genes - showed such enrichment (**Fig. 4e**): The Turquoise and Brown modules were enriched for XCI cluster genes, whereas the Green and Blue modules were enriched for XCIE cluster genes. The Blue module was unique for its additional enrichment in PAR genes, and its inclusion of XIST. We generated network visualizations to more closely examine SCD-sensitive genes and gene co-expression relationships within each of these four sex-chromosome enriched WGCNA modules (**Fig. 4f** Blue and **Fig. S3c-e** for others, **Methods)**. The Blue module network highlights XIST, select PAR genes (*SLC25A6*, *SFRS17A*) and multiple X-linked genes from X-Y gametolog pairs (*EIF1AX*, *KDM6A* (*UTX*), *ZFX*, *PRKX*) for their high SCD-sensitivity, and shows that these genes are closely co-expressed with multiple SCD-sensitive autosomal genes including *ZWINT*, *TERF2IP* and *CDKN2AIP*.

Our detection of highly-organized co-expression relationships between SCD sensitive sex-linked and autosomal genes hints at specific regulatory effects of dosage sensitive sex chromosome genes in mediating the genome-wide effects of SCD variation. To test this, and elucidate potential regulatory mechanisms, we performed an unbiased transcription factor binding site (TFBS) enrichment analysis of genes within Blue, Green, Turquoise and Brown WGCNA modules (**Methods**). This analysis converged on a single TF - ZFX, encoded by the X-linked member of an X-Y gametolog pair – as the only SCD sensitive TF showing significant TFBS enrichment in one or more modules. Remarkably, the gene *ZFX* was itself part of the Blue module, and ZFX binding sites were not only enriched amongst Blue and Green module genes (increased in expression with increasing X-chromosome dose), but also amongst Brown module genes that are downregulated as X-chromosome dose increases (**Fig. 4g)**. To directly test if changes in ZFX expression are sufficient to modify expression of Blue, Green or Brown modules genes in immortalized lymphocytes, we harnessed existing gene-expression data from murine T-lymphoblastic leukemia cells with and without ZFX knockout (19) (GEO GSE43020). These data revealed that genes downregulated by ZFX knockout in mice have human homologs that are specifically and significantly over-represented in Blue (p=0.0005) and Green (p=0.005) modules (p>0.1 for each of the other 6 WGCNA modules) – providing experimental validation of our hypothesized regulatory role for ZFX.

## DISCUSSION

In conclusion, our study – which systematically examined gene expression data from 470 individuals representing a total of 9 different sex chromosome karyotypes - yields several new insights into sex chromosome biology with consequences for basic and clinical science. First, our discovery and validation of X-linked genes that are upregulated by reducing X-chromosome count – so that their expression is elevated in XO vs. XX for example – runs counter to dominant models of sex chromosome dosage compensation in mammals, and thereby modifies current thinking regarding which subsets of X-chromosome genes could contribute to sex and SCA-biased phenotypes (20). We speculate that this newly-observed non-canonical mode of X-chromosome dosage sensitivity could arise through one or more of the following candidate mechanisms: (i) repression of X-linked genes on the active X-chromosome by genes that are expressed from inactive X-chromosomes, and (ii) sensitivity of X-linked genes on the active X-chromosome to the changes in nuclear heterochromatin dosage that would follow from varying numbers of inactivated X-chromosomes (21).

Our findings also modify classical models of sex-chromosome biology by identifying X-linked genes that vary in their expression as a function of Y–chromosome dosage – indicating that the phenotypic effects of normative and aneuploidic variations in Y-chromosome dose could theoretically be mediated by altered expression of X-linked genes. Moreover, the discovery of Y chromosome dosage effects on X-linked gene expression provides novel routes for competition between maternally and paternally inherited genes beyond the previously described mechanisms of parental imprinting and genomic conflict–with consequences for our mechanistic understanding of sex-biased evolution and disease (22).

Beyond their theoretical implications, our data help to pinpoint specific genes that are likely to play key roles in mediating sex chromosome dosage effects on wider genome function. Specifically, we establish that a distinctive group of sex-linked genes - notable for their evolutionary preservation as X-Y gametolog pairs across multiple species, and the breadth of their tissue expression in humans (10) – are further distinguished from other sex-linked genes by their exquisite sensitivity to SCD, and exceptionally close co-expression with SCD-sensitive autosomal genes. These results add critical evidence in support of the idea that X-Y gametologs play a key role in mediating SCD effects on wider genome function. In convergent support of this idea we show that (i) multiple SCD-sensitive modules of co-expressed autosomal genes are enriched with TFBS for an X-linked TF from the highly dosage sensitive *ZFX-ZFY* gemetolog pair, and (ii) ZFX deletion causes targeted gene expression changes in such modules. Inclusion of ZFX in a co-expression module (Blue) with enriched annotations for chromatin organization and cell cycle pathways is especially striking given the rich bodies of experimental data which have independently identified ZFX as a key regulator of cellular renewal and maintenance (23).

Gene co-expression analysis also reveal the diverse domains of cellular function that are sensitive to SCD – spanning cell cycle regulation, protein trafficking and energy metabolism. These effects appear to be specific to shifts in SCD as they are not induced by trisomy of chromosome 21. Furthermore, gene co-expression analysis of SCD effects dissects out specific immune activation pathways that are upregulated by supernumerary X-chromosomes in males, and enriched for genes known to confer risk for autoimmune disorders that are overrepresented amongst males bearing an extra X-chromosome. Thus, we report coordinated genomic response to SCD that could potentially explain observed patterns of disease risk in SCA.

Collectively, these novel insights serve to refine current models of sex-chromosome biology, and advance our understanding of genomic pathways through which sex chromosomes can shape phenotypic variation in health, and sex chromosome aneuploidy.

## MATERIALS AND METHODS

### Acquisition and preparation of biosamples

RNA was extracted by standard methods (Qiagen, MD, USA) from lymphoblastoid cell lines (LCLs) for 469 participants recruited through studies of SCA at the National Institute of Health Intramural Research Program, and Thomas Jefferson University (24). All participants with X/Y-aneuploidy were non-mosaic, and stability of karyotype across LCL derivation was confirmed by chromosome fluorescent in situ hybridization (FISH) in all members of a randomly selected subset of 9 LCL samples representing each of the 4 supernumerary SCA groups included in our microarray analysis. Sixty-eight participants provided RNA samples for microarray analyses (12 XO, 10 XX, 9 XXX, 10 XY, 8 XXY, 10 XYY, 9 XXYY), and 40 participants provided RNA samples for a separate qPCR validation/extension study (4 XO, 145 XX, 22 XXX, 146 XY, 34 XXY, 16 XYY, 17 XXYY, 8 XXXY, 10 XXXXY). The microarray and qPCR samples were fully independent of each other (biological replicates), with the exception of 2 XO participants in the microarray study, who each also provided a separate LCL sample for the qPCR study (**Table S1**).

### Microarray data preparation, differential expression analysis, annotation and probe selection

Gene expression was profiled using the Illumina HT-12 v4 Expression BeadChip Kit (Illumina Inc, San Diego, CA). Expression data were quantile normalized across arrays and log2 transformed using the *limma* package in R(25). For each of 47323 probes, we estimated mean expression by karyotype group, and log2 fold change in gene expression for each unique pairwise group contrast between karyotype groups (**Fig. 1a**), along with their associated false-discover-rate (FDR) corrected p-values. For each pairwise karyotype group comparison, we identified probes with significant log2 fold-changes that survived FDR correction for multiple comparisons across all 47323 probes with q (the expected proportion of falsely rejected nulls) set at 0.05. We also calculated a single summary estimate of SCD effects for each probe, by calculating the proportion of variance (r^2^) in probe expression that was accounted for by the 7-level factor of SCD group.

All 47323 microarray probes were annotated using both the vendor manifest file and an independently published re-annotation that assigns a quality rating to each probe based on the specificity of its alignment to the purported transcript target (26). We filtered for all probes with “perfect” or “good” quality alignment to a known gene according to this reannotation, and then used the *collapseRows* function from the WGCNA (27) package in R (with default settings), to select one probe per gene. We also applied a further filter to remove any Y-linked probes that showed differential expression between female karyotype groups. These steps resulted in high-quality measures of expression and estimates of differential expression for 19984 autosomal and 894 sex-chromosome genes in each of 68 independent samples from 7 different karyotype groups.

To select a log2 threshold for use in categorical definition of differentially expressed genes (DEGs), we estimated DEG count across a range of absolute log2 fold-change cut-offs for 6 contrasts of primary interest: karyotypically normal males vs. females (i.e. XY vs. XX), and each SCA group vs. its respective “gonadal control” (i.e XX was the control for XO and XXX groups / XY was the control for XXY, XYY and XXYY). An absolute log2 fold change of 0.26 (equivalent to a ~20% increase/decrease in expression) was empirically selected as the cut-off to define differential expression by (i) separately identifying the increase in log2FC threshold that cause the greatest drop in DEG count for each SCA group, (ii) averaging these log2FC thresholds across all 5 SCA groups. Thus in any contrast between two karyotype groups, DEGs showed log 2 fold-changes that met both of the following two criteria: an associated p value that met FDR correction with q<0.05, and an absolute magnitude greater than 0.26.

### Identifying Genes Showing Significant Differential Expression Across All Sampled Changes in X- and Y-Chromosome Count

The 21 unique pairwise karyotype group comparisons in our microarray dataset included 15 contrasts involving a disparity in X-chromosome count, and 16 contrasts involving a disparity in Y-chromosome count (**Fig. 1a**). Using the empirically-defined |log2 fold change| cut-off of 0.26 (see above), we screened all 20878 genes for evidence of significant differential expression across all instances of X- or Y-chromosome disparity.

### Comparing SCD Sensitivity of Gametolog vs. non-Gametolog Sex-Linked Genes

Fourteen of 16 established (9) X-Y gametolog gene pairs were represented in our microarray dataset. To compare the SCD sensitivity of this gene set to that of non-gametolog sex-linked genes we first quantified the mean effect of SCD group on gene expression within the gametolog gene set by averaging gene-wise r-squared values for the effect of SCD group on expression. We then determined the centile of this observed gene set mean r-squared against a distribution of r-squared values for 10,000 similarly sized sets of randomly sampled non-gametolog sex-linked genes (**Fig. 1d**). This procedure was conducted separately for X- and Y-chromosomes.

### A Priori Assignment of Sex Chromosome Genes to Four Class Model Categories

PAR genes were defined as those lying distal to the PAR1 and PAR2 boundaries specified in hg18 build of the human genome. Y-linked and X-linked genes were defined as those lying proximal to these PAR boundaries on the Y- and X-chromosome respectively. X-linked genes were assigned to XCIE and XCI using consensus classifications from a systematic integration (8) of XCI calls from 3 large-scale assays: expression from the inactivated human X-chromosome in 9 hybrid human-mouse cell lines (12), allelic-imbalance analysis of expression data in cell-lines from females with skewed X-inactivation (28), and X-chromosome methylation data from microarray (29). According to the XCI categories of this consensus report we classified X-linked genes as X-inactivated (“Subject” or “Mostly Subject” categories), X-escape (“Escape” or “Mostly Escape” categories), or X-other (all other intermediate categories).

### Clustering of Sex Chromosome Genes by Dosage Sensitivity

For all 894 sex chromosome genes within our dataset we calculated the mean fold-change per SCD group relative to mean expression across all SCD groups. The resulting 894 by 7 matrix was submitted to k-means clustering across a range of k-values using the *kmeans* function in R with nstart and iter.max set at 100. Visual inspection of a scree plot of mean within partition sum-of-squared residuals against k indicated an optimal 6-cluster solution (**Fig. S2a**). The largest of these 6 clusters (Gray cluster) gathered genes with low or undetectable expression levels across all samples, and was excluded from further analysis.

Reproducibility of 6 cluster solution was established using a bootstrap method whereby individuals were randomly drawn (with replacement) from each SCD group within our microarray dataset to derive 1000 bootstrap sets of 68 samples. k-means clustering was repeated for each of these 1000 sets to define a 6-cluster solution in each draw (**Fig. S2b**). Consistency of clustering was quantified for each gene as the proportion of bootstrap draws in which it was assigned to the same cluster as it had been in the original sample. The median consistency score for cluster designation was >93% for all 5 clusters of SCD-sensitive sex chromosome genes.

The observed grouping of sex chromosome genes from k-mean clustering, was compared with the predicted Four Class Model groupings using two-tailed Fishers tests for all pairwise cluster-grouping combinations (**Fig. 2b**).

### Modelling X- and Y-chromosome Dosage Effects on Expression of Sex Chromosome Gene Clusters

We used the following linear models to estimate the combined influence of X and Y chromosome dosage in cluster and gene-level expression of Yellow (XCI enriched) and Green (XCIE enriched) gene clusters:

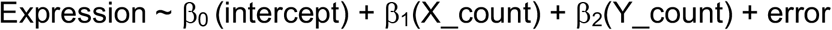

We computed p-values for comparisons of both β_1_ and β_2_ coefficient estimates against the null (0), and used these to test for significant directional effects of sex chromosome dosage on the mean expression of each gene cluster, as well the expression of individual genes within each cluster.

### Aligning Sex Chromosome Expression, Epigenetic and Evolutionary Data

We validated our data-driven clustering of X-linked genes into Yellow (XCI enriched) and Green (XCIE enriched) groups, by overlapping the genomic coordinates of gene probes with segmentations of the X-chromosome according to (i) “chromatin states” defined by computational analysis of coordinated changes in 10 distinct chromatin marks in LCLs (16), (ii) “evolutionary strata” reflecting staged loss of recombination between the X- and Y-chromosome (15). Overlaps of our probe coordinate with these two annotations were defined using the GenomicRanges package in R. As a third validation we also aligned our clustering of X-lined genes with a previously published annotation of X-linked genes according to whether their corresponding ancestral Y-linked homologue has been lost, converted to a pseudogene or maintained (8). Non-random associations between these three annotations and Yellow vs. Green k-means cluster membership were assessed using Chi-squared tests.

### Comparison of DEG Count and Genomic Distribution Across SCA Groups

Total DEG counts were compared across SCD groups across a range of log 2 fold change cut-offs as described above and reported in **Fig. 3 a,b**. To test for non-random distribution of DEGs across the genome in each SCD group **(Fig. 3c**), we compared observed DEG counts across 4 genomic regions - autosomal, PAR, Y-linked and X-linked - to the background distribution of total gene counts across these regions using the prop.test function in R. All SCD groups showed a high non-random distribution of DEGs across the genome – reflecting preferential involvement of sex chromosome genes (p < 7.2 * 10^−13^).

### Quantitative rtPCR validation of Differentially-Expressed Genes in Microarray

#### Selecting genes of interest

For selected genes showing significant differential expression between karyotype groups in our core sample, we used qPCR to validate and extent observed fold-changes in an independent sample of 402 participants representing all the karyotypes in our core sample plus two additional SCAs: XXXY, and XXXXY. Selection of specific genes for qPCR validation was as follows. From the set of 10 sex-linked genes with patterns of “obligate” dosage sensitivity (**Fig 1b**), we selected XIST, the 2 most X-chromosome sensitive X-linked gametologs [EIF1AX, KDM6A(UTX)], and the 2 most Y-chromosome sensitive Y-linked gametologs (ZFY, DDX3Y). All genes selected for qPCR from the sets of dosage sensitive sex chromosome genes defined by k-means clustering (**Fig. 2a,b**) showed (i) stable cluster membership in >95% of bootstrap draws (**Fig. S2b**), and (ii) consistent inclusion in the top 10 DEGs across multiple relevant group contrasts for that k-means cluster (Pink Y-linked cluster: XYY vs. XY and XXYY vs. XXY | Yellow XCI and Green XCIE clusters: XO vs. XXX, XO vs. XY, XXY vs. XX).

#### Fluidigm qPCR protocol

Reverse Transcription reaction was performed using RT2 HT First Strand Kit (QIAGEN, 330411) with 1000 ng RNA input per sample. One-tenth of cDNA was preamplified using RT2 Microfluidics qPCR Reagent system (QIAGEN, 330431) in combination with custom RT2 PreAmp pathway primer mix Format containing 94 RT2 primer assays. Fourteen cycles of preamplification were performed using the manufacturer recommended preamplification protocol. Amplified cDNA was diluted 5-fold using RNase-free water and assessed in real-time PCR using the RT2 Micrifluidics EvaGreen qPCR Master mix and a Custom RT2 profiler PCR array PCR Array containing 96 assays, including selected DEGs of interest, housekeeping genes, reverse-transcription controls, and positive PCR control. Real-time PCR was performed on a Fluidigm BioMark HD (Fluidigm, San Francisco, US) using the RT2 cycling program for the Fluidigm BioMark, which consists of an initial thermal mix stage (50°C for 2 minutes, 70°C for 30 minutes, and 25°C for 10 minutes) followed by a hot start at 95°C for 10 minutes and 40 cycles of 94°C for 15 seconds, and 60°C for 60 seconds. For data processing, an assay with Ct > 23 was deemed to be not expressed.

#### Differential Expression Analysis of qPCR data

The ΔΔCT method of relative quantification was used to analyze qPCR data (30). To provide normalized estimates of expression for each gene we calculated ΔCT values, by subtracting the CT for each gene of interest from the mean CT of two housekeeping genes (GAPDH and RPLP0) which were not differentially expressed across groups in either microarray or qPCR data. Thus, larger ΔCT values reflected greater normalized expression relative to mean expression of the reference housekeeping genes. These ΔCT values were used as input for calculation of all unique pairwise group differences in expression between karyotype groups represented in the independent qPCR validation dataset. Group differences in expression were modeled using the limma R package with identical setting to those used in analysis of microarray data (see above). The resulting ΔΔCT represent fold-changes in gene expression between groups, on a log scale with a base determined by the effective qPCR efficiency.

#### Validation Microarray Results Using qPCR Results

All 21 unique pairwise SCD group contrasts in our microarray sample could be reproduced in the independent qPCR dataset. We used the correlation across these 21 group contrasts for the qPCR fold-change and microarray log2 fold change to quantify the degree of agreement between qPCR and microarray findings (**Fig. S1a and Fig. S2d,e**).

#### Extension of Microarray Results Using qPCR Results

The qPCR dataset also included two SCD groups that were not represented in the microarray dataset – XXXY and XXXXY – allowing for a total of 15 novel pairwise SCD group contrasts (“XO-XXXY”, “XO-XXXXY”, “XXXY-XX”, “XXXXY-XX”, “XXX-XXXY”, “XXX-XXXXY”, “XXXY-XY”,“XXXXY-XY”, “XXY-XXXY”, “XXY-XXXXY”, “XYY-XXXY”, “XYY-XXXXY”, “XXYY-XXXY”, “XXYY-XXXXY”, “XXXY-XXXXY”) sampling diverse disparities of X- and Y-chromosome dosage. These novel contrasts were used as a further test for the validity and reproducibility of our microarray findings. Each of the 15 novel pairwise SCD group contrasts was coded according to two effects of interest: difference in X-chromosome count and difference in Y-chromosome count. These coded SCD disparities were then correlated with observed fold-changes for unique pairwise group contrasts in the qPCR dataset to test if patterns of fold-change observed in the microarray dataset could be extension into unseen karyotype groups (**Fig. S1b**,**c**: X- and Y-linked genes with “obligatory” sex chromosome dosage sensitivity | **Fig. S2f,g**: X-linked genes from the “Yellow” and “Green” k-means that countered expectations of the classical Four Class Model).

### Weighted Gene Co-expression Network Analysis (WGCNA)

#### Defining Gene Co-expression Modules

Gene co-expression modules were generated using the R package Weighted Gene Co-expression Network Analysis (WGCNA). Briefly, this involved first calculating the Pearson correlation coefficient between all 20978 genes across all 68 samples in our study. This correlation matrix was transformed using a signed power adjacency function with a threshold power of 12 (selected based on fit to scale-free topology), and then converted into Topological Overlap Matrix (TOM) by modifying the correlation between each pair of genes using a measure of the similarity in their respective correlations with all other genes (31). The resulting TOM was then converted to a distance matrix by subtraction from 1, and used to generate a dendrogram for clustering genes into modules. Gene modules were defined using the Dynamic Hybrid cutree function (32) [with the following parameter settings: deepSplit (control over sensitivity of module detection to module splitting) = 2, mergeCutHeight (distance below which modules are merged)= 0.25, minimum module size=30)]. Given the large number of genes included in our analyses, we implemented module detection using the “blockwise” WGCNA algorithm, which starts with a computationally inexpensive method to assort genes into smaller co-expression blocks, and then completes the above steps within each block before merging module designations across blocks. This implementation of WGCNA defined 18 mutually exclusive co-expression gene modules within our data, which ranged from 45 to 1393 genes in size, and a left-over group of 14630 genes without module membership (**Table S3**). The expression of each module was summarized as a module eigengene value (ME: the right singular vector of standardized expression values for genes in that module) in every sample. These ME values were used to determine differential expression of modules across (omnibus F-tests) and between (T-tests) SCD groups, as well as module co-expression across samples (Pearson correlation coefficient).

#### Further characterizing gene co-expression modules

We used module preservation analysis to establish that our defined co-expression modules were not dominated by (i) the large number of DEGs induced by X-monosomy (using expression data excluding XO samples), or (ii) other SCD group differences in mean expression levels (using expression data after residualization for the effects of SCD group and re-centering at a common mean). All modules showed high reproducibility based on a module-specific Z_summary_ scores derived by comparing observed modular connectivity and density metrics with null values generated by 200 permutations of gene-level module membership(33). We focused further characterization of modules which passed two independent statistical criteria; (i) SCD sensitivity - quantified using F-tests for the omnibus effects of karyotype group on modular expression quantified as the ME, (ii) functional coherence as inferred by analysis of modular gene ontolology term enrichments using GO elite (34), and Gorilla (35).

#### Testing for enrichment of autoimmune disorder risk genes in WGCNA modules

A large-scale records-based study was used to define 10 Autoimmune Disorder (ADs) with clearly elevated prevalence rates in XXY vs. XY males (7), 9 of which were represented in the largest available catalog of Genome Wide Association Study (GWAS) findings (https://www.ebi.ac.uk/gwas/): Diabetes Mellitus type 1, Multiple Sclerosis, Autoimmune Hypothyroidism, Psoriasis, Rheumatoid Arthritis, Sjogren’s Syndrome, Systemic Lupus Erythematosus, Ulcerative Colitis, and Coeliac Disease. A total of 495 genes within our microarray sample were annotated for showing a significant association in GWAS with one or more of these 9 AD conditions. Overrepresentation of this AD gene set in the XXY upregulated Red gene co-expression module was tested for using both Fisher’s exact test (p=0.01), and by comparing the observed representation of AD genes against a null distribution generated by 10,000 random gene samples of equal size to the red module.

#### Testing for patterned enrichment of dosage sensitive sex chromosome genes in WGCNA modules

We tested if any of the 8 SCD-sensitive and functionally enriched WGCNA modules showed enrichment for the previously derived k-means clusters of dosage sensitive sex chromosome genes (**Fig. 2**) by applying two-tailed Fishers tests to all pairwise module-cluster combinations (**Fig. 4e**). All observed associations survived Bonferroni correction for multiple comparisons.

#### Module Visualization

WGCNA co-expression modules were visualized by selecting genes within the top decile of SCD-sensitivity (indexed using r-squared for proportion of expression variance explained by group), and edges (co-expression links between genes) in the top decile of edge strengths. All visualizations were constructed using the igraph R package in R.

#### Transcription factor binding site analyses

Transcription factor binding site (TFBS) enrichment analysis was performed each of the 4 SCD-sensitive WGCNA modules - Blue, Green, Turquoise and Brown - that were enriched for inclusion of gene from one or more of the 5 SCD-sensitive clusters of sex chromosome genes. In each module, we scanned canonical promoter regions (1000bp upstream of the transcription start site) for the top 500 genes with strongest intramodular “connectivity” (based on kME - the magnitude of each gene’s coexpression with its module’s eigengene). Next we utilized TFBS position weight matrices (PWMs) from JASPAR database (205 non-redundant and experimentally defined motifs) (36) to examine the enrichment for corresponding TFBS within each module. For TFBS enrichment all the modules were scanned with each PWMs using the Clover algorithm (37). To compute the enrichment analysis, we utilized three different background datasets (1000 bp sequences upstream of all human genes, human CpG islands and human chromosome 20 sequence). To increase confidence in the enrichment analyses, we considered TFBS to be over-represented based on the P-values (<0.05) obtained relative to all the three corresponding background datasets.

#### Enrichment of SCD sensitive modules for genes with DE due to experimental ZFX knockout

To provide an orthogonal experimental test for evidence of a regulatory role for ZFX within Blue, Green or Brown WGCNA modules, we used a list of genes with significantly decreased expression due to ZFX knockout in murine lymphocytes(19). Human homologs were found for these genes (<u>http://www.informatics.jax.org/downloads/reports/index.html</u>), and two-tailed Fishers Tests were used to assess if these human genes were significantly enriched/impoverished in any of the 8 SCD sensitive WGCNA modules.

### Comparison of Autosomal Gene Fold-Change in SCA and Down Syndrome

The transcriptomic effects on Trisomy 21 (T21) were characterized in LCLs by passing a publically available Illumina microarray gene expression dataset (GEO, Accession number GSE34458) through an identical analytic pipeline to that used in characterizing genome-wide fold changes in our SCA sample (see above). We first independently confirmed the previously reported finding that chromosome 21 was robustly enriched for genes showing differential expression in this T21 data set (Chi-squared=999, p<2*10-16 for enrichment of DEGs on chromosome 21) – buttressing use of these data to assess transcriptomic effects of T21. We examined overlaps in genome-wide expression change between T21 and the three sex-chromosome trisomes in our samples (XXY, XYY and XXX) using 17671 genes with complete expression data in both microarray datasets (after exclusion of genes on chromosomes X, Y and 21). We tested for, and failed to find any evidence of significant overlap in DEGs using Chi-squared tests (**Table S4**). To test if T21 showed a similar shift in gene co-expression modules to sex chromosome trisomies, we used the designation of genes to modules in the SCA sample to recalculate module Eigengenes. We then calculated MR fold changes for T21 and three SCA trisomies (XXX, XXY and XYY). Trisomy of chromosome 21 was associated with a clearly distinct profile of ME expression chance than all three of the SCA trisomies (**Fig. 4c**).

